# Virus survival in evaporated saliva microdroplets deposited on inanimate surfaces

**DOI:** 10.1101/2020.06.15.152983

**Authors:** Aliza Fedorenko, Maor Grinberg, Tomer Orevi, Nadav Kashtan

## Abstract

The novel coronavirus respiratory syndrome (COVID-19) has now spread worldwide. The relative contribution of viral transmission via fomites is still unclear. SARS-CoV-2 has been shown to survive on inanimate surfaces for several days, yet the factors that determine its survival on surfaces are not well understood. Here we combine microscopy imaging with virus viability assays to study survival of three bacteriophages suggested as good models for human respiratory pathogens: the enveloped Phi6 (a surrogate for SARS-CoV-2), and the non-enveloped PhiX174 and MS2. We measured virus viability in human saliva microdroplets, SM buffer, and water following deposition on glass surfaces at various relative humidities (RH). Although saliva microdroplets dried out rapidly at all tested RH levels (unlike SM that remained hydrated at RH ≥ 57%), survival of all three viruses in dry saliva microdroplets was significantly higher than in water or SM. Thus, RH and hydration conditions are not sufficient to explain virus survival, indicating that the suspended medium, and association with saliva components in particular, likely affect physicochemical properties that determine virus survival. The observed high virus survival in dry saliva deposited on surfaces, under a wide range of RH levels, can have profound implications for human public health, specifically the COVID-19 pandemic.

## Introduction

The novel human coronavirus that emerged in China in late 2019 caused a pandemic. Human saliva microdroplets expelled into the air through coughing, talking, and breathing are considered a key source of transmission of the virus ^1,2^. These microdroplets, ranging in size between few micrometers up to millimeters ^3–5^ travel in the air, while some of them – larger ones in particular – settle on surfaces ^2^. Thus, inanimate surfaces are a potential medium of virus transmission ^1,2,6,7.^ Like many other viruses, SARS-CoV-2 has been shown to survive well and remain viable on various surfaces, e.g., metal, glass, and plastic, for up to several days ^8–10^. While not necessarily viable, SARS-CoV-2 RNA has been detected on surfaces in contaminated sites such as hospitals ^11–14^.

The factors that affect virus survival in droplets that settle on surfaces are complex and not well understood. Survival is expected to depend upon the physicochemical characteristics and hydration conditions of the immediate microscopic environment of the virion. These in turn are determined by several factors, including the composition of the fluid comprising the droplets, the surface properties, the ambient temperature, and the relative humidity (RH). A preeminent source for expelled droplets is human saliva, a complex solution that contains salts, a variety of proteins, and surfactants ^15–18^. It has been suggested that micrometer-sized dry deposits of saliva droplets on surfaces protect virions ^19^. Still, it is not clear how well viruses survive in saliva microdroplets following their deposition on surfaces, and what factors are in play therein. The impact of RH on the stability and viability of SARS-CoV-2 and other enveloped viruses has been studied mostly in airborne droplets or aerosols ^20–26^ and less on surfaces ^27–29^.

While survival varies between virus species, increased survival at both low (<40%) and very high (>90%) RH is often observed, with decreased survival at intermediate RH levels ^20–22,25,27.^ The underlying mechanism of this U-shaped survival as a function of RH is not clear ^24^. Only a few studies have attempted to gain mechanistic understanding of how RH affects virus stability in microdroplets. Prominent among these are the pioneering studies of the Marr group ^5,24,26,30,^ which point to the role of the solvent composition, evaporation dynamics, and RH on virus survival in aerosols and sessile droplets.

A key factor determining the evaporation rate and equilibrium hydration level of drying droplets on surfaces (and in air as well) is the deliquescence property of solutes, or mostly highly hygroscopic salts ^31–33^. Accordingly, although many surfaces in the indoor environment appear dry, they are in fact covered by thin liquid films and micrometer-sized droplets, invisible to the naked eye, known as *microscopic surface wetness*, or MSW ^34^. MSW can be considered the “envelope” that accommodates microorganisms on surfaces, and as such has profound impact on many aspects of microbial life. For example, it can protect microbes from complete desiccation. However, MSW is a harsh micro-environment that differs in its properties from bulk liquid. MSW inherently arises from drying liquids that evaporate on a surface. This drying process is accompanied by physicochemical changes such as solute concentrations, pH, reactive oxygenic species, surface tension, and others. At the microscale, gradients, local densities, and surface irregularity introduce heterogeneity to MSW environments ^18,35–37^. Collectively, MSW imposes severe stresses on microbes therein – including viruses – and affects their survival ^18,26,27,38,39^.

The current study, motivated by the urgent call for the scientific community to address SARS-CoV-2 spread, aims to explore the link between microscopic surface wetness and virus survival therein. We focus on two variables that directly affect MSW – solution composition and RH levels – and seek to determine whether and how it is reflected by virus survival trends. We used bacteriophages suggested as good, safe, and easy to work with, model surrogates to study surface and air survival of pathogenic viruses ^40–42^. We chose to study three phages: the enveloped Phi6 – proposed as a good surrogate for SARS coronaviruses – and two other tailless non-enveloped model bacteriophages as a reference. Phi6 is a dsRNA phage of the *Cystoviridae* family that has been suggested as a good surrogate for studying enveloped RNA viruses ^18,22,40,43–45^; similar to SARS-CoV-2, it is enveloped by a lipid membrane, has spike proteins, and is of similar size (~80-100 nm). The other virus strains we used are the well-studied, non-enveloped MS2 (ssRNA; *Leviviridae*) ^41,46–48^ and PhiX174 (ssDNA; *Microviridae*) ^41,49^. To better understand how RH and the solution composition of microdroplets affect virus survival on surfaces, we combine microscopy imaging to assess MSW state with plaque assays to determine virus infectivity. We compare survival in ‘sprayed’ microdroplets of three suspensions: human saliva, SM buffer, and ‘pure’ water under a range of RH (23%-78%) relevant to the indoor environment. The link between the microscopic environment of viruses, including hydration conditions, and virus viability is discussed.

## Materials and Methods

### Bacteria and bacteriophage strains

Bacteriophages Phi6 (DSM-21518), PhiX174 (DSM-4497), and MS2 (DSM-13767) were purchased from DSMZ. Vacuum-dried phages were revitalized according to DSMZ instructions. Bacteriophage propagation: Overnight bacterial host cultures were diluted at 1:50 into 1 ml fresh media and grown to OD_600_ 0.3. A single plaque, suspended in 100 μl SM buffer, was used to inoculate the fresh culture. The first propagation was shake incubated at the appropriate temperature for the host strain for 3 hrs, or until lysis was observed. Meanwhile, the overnight cultures were diluted again in 9 ml fresh media and grown to OD_600_ 0.3. The first propagation was then diluted with the 9 ml fresh host cultures and grown for 3 hrs or until lysis was observed. The second propagation was centrifuged (4,200_RCF_, 10 minutes) and filtered through a 0.22 μm filter (Millex® GV). Phages were stored at 4°C and their concentrations were determined by the drop plaque assay. *Pseudomonas syringae* (DSM-21482), a host strain for bacteriophage Phi6, was purchased from the German Collection of Microorganisms and Cell Cultures (DSMZ) and cultivated in TSB (Tryptic Soy Broth) at 28°C. *Escherichia coli* (DSM-13127), a host strain for bacteriophage PhiX174, was purchased from the German Collection of Microorganisms and Cell Cultures (DSMZ) and cultivated in LB (Luria Bertani broth) at 37°C. *Escherichia coli* (ATCC® 15597™) (kindly provided by Adi Stern), a host strain for bacteriophage MS2, was cultivated in LB (Luria Bertani broth) at 37°C.

### Viability of bacteriophages in microdroplets deposited on a glass surface

Phage stock solution (10^8^ PFU/ml for Phi6; and MS2 and 10^7^ PFU/ml for PhiX174) were diluted 100fold with: (1) SM Buffer (100 mM NaCl, 8mM MgSO4×7H_2_O, 50 mM Tris-Cl pH 7.5, 0.01% w/v gelatin) (2) filter-sterilized H_2_O (W4502, sigma) and (3) natural human saliva (donated by one of the authors). Fluorescent beads (2 μm, melamine resin labeled by rhodamine B, 94009 Fluka) were added to the suspension with a 5×10^2^ final dilution. Solutions were loaded into 5ml refillable spray bottles (purchased at a local cosmetics store) and a portion of the load was sprayed on a 12-well glass bottom plate (P12-1.5H-N, Cellvis). Each well was sprayed by pressing the spray nozzle twice, delivering a volume of ~50 μl). Spray was applied to the well through a 50 ml falcon tube from which we chopped 1.5 cm from its conical end. The plates were placed without the cover lid inside a sealed plastic box. A 100 ml saturated salt solution was placed in the bottom of each box to maintain relative humidity of 23, 43, 57, and 78% (potassium acetate, potassium carbonate, sodium bromide, and ammonium chloride respectively). The boxes were placed in an incubator set at 25°C for 14 hrs. Control: spray bottles with the remaining (unsprayed) suspensions were left in incubation for 14 hrs and sprayed onto the 12-well plates at the end of the incubation period. All plates were imaged at the end of the 14-hr incubation period and then suspended with 500 μl of SM buffer for the drop plaque assay.

### Virus Plaque Assay

Plates containing the bottom agar layer were poured in advance (TSB or LB with 15g/l agar, 5 mM MgSO_4_, and 5 mM CaCl_2_). On the day of the experiment, overnight bacterial host cultures were diluted at 1:50 into 1 ml fresh media and shake incubated until they reached OD_600_ 0.3. Meanwhile, the top agar (TSB or LB with 7.5g/l agar, 5 mM MgSO_4_, and 5 mM CaCl_2_) was melted and kept in a 55°C water bath. The bacterial culture was combined with the top agar at a ratio of 1:40, and poured on top of the bottom layer. The phages were serially diluted in SM buffer, and after the top agar solidified, either 10 μl, 100 μl, or 1000 μl were pipetted and spread out onto either one quarter, one half, or a full plate (10×10 cm petri dish) respectively. Plates were left open until dry and incubated at the appropriate temperature for the host strain (as described above) overnight.

### Microscopy

12-well plates were mounted on a stage top without warming (room temperature) during image acquisition. Images were collected using an Eclipse Ti-E inverted epi-fluorescence microscope (Nikon, Japan) equipped with a Plan Apo 20x/0.75 NA air objective and the Perfect Focus System for maintenance of focus. An LED light source (SOLA SE II, Lumencor) was used for fluorescence excitation. Rhodamine B-marked fluorescent beads were excited through a 560/40 filter, and emission was collected with a T585lpxr dichroic mirror and 630/75 filter (filter cube #49008, Chroma). Images were acquired with a SCMOS camera (ZYLA 4.2PLUS, Andor Technology Ltd., UK) controlled with NIS Elements 5.02 software (Nikon Instruments Inc., USA). 10×10 adjacent fields of views (covering a total area of 2.82 × 2.82 mm) were monitored per each well. Multiple stage positions were collected using a motorized encoded scanning stage (SCANplus IM 130 × 85, Märzhäuse).

## Results

To study virus survival in microdroplets deposited on a smooth inanimate surface, we sprayed Phi6, MS2, and PhiX174 viruses suspended in three media – human saliva, water, and SM buffer – on glass-bottom 12-well plates (Fig. 1, Methods). Fluorescent beads (2 μm in diameter) in an equivalent concentration to those of the Phi6 and MS2 viruses (~10^6^/ml), were added to the suspension for two purposes: (i) to mimic micrometer-sized particulates (e.g., bacteria cells) that are spread on real-world surfaces and have been shown to affect the formation of microscopic surface wetness ^34^; and to (ii) help interpret the virion distribution within droplets. Sprayed microdroplet size ranged between tens to hundreds of microns in diameter (Fig. 2 A,B), which falls within the range of respiratory fluid microdroplets exhaled while coughing, speaking, and breathing ^4,5^ and gravitate toward surfaces (i.e., not the < 5 μm aerosols). As this study’s emphasis is virus survival in the indoor environment, we chose to work at a temperature of 25°C and a range of RH (23% to 78%) that spans most indoor environments ^50^. The sprayed well plates were placed under constant temperature and RH conditions for 14 hrs, and subsequently microscopy images were taken and virus survival was estimated by the plaque assay using the corresponding bacterial host as a reporter for infectivity (Fig. 1, Methods).

**Fig. 1.**
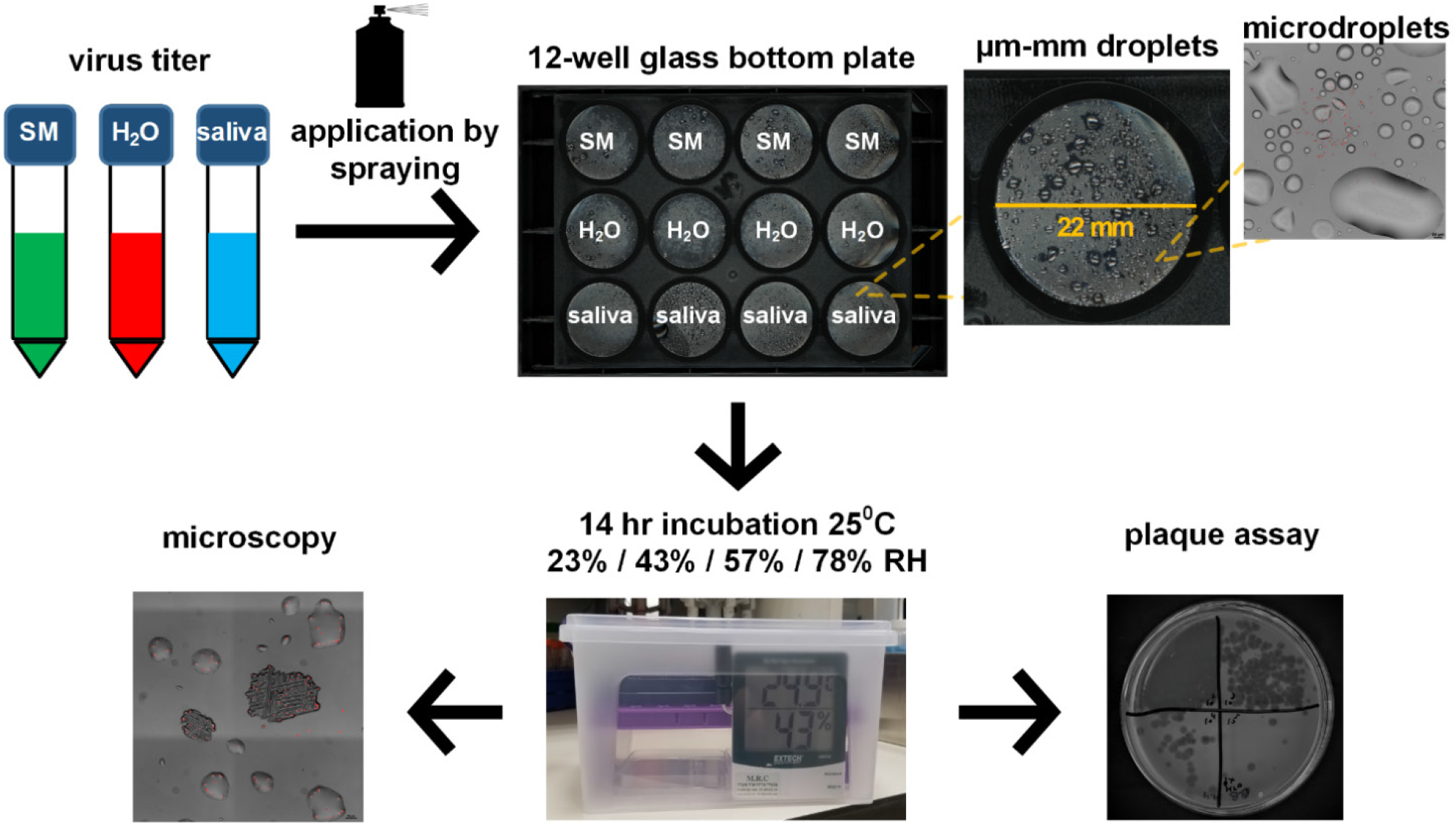
Schematic illustration of the experimental workflow. Phage suspended in SM buffer, H_2_O, and human saliva were applied onto a glass-bottom 12-well plate by a spraying device, resulting in microdroplets ranging in size from ~10 microns to ~1 mms in diameter. Plates were transferred to sealed containers that maintained specific RH conditions (using saturated salt solutions), and the containers were placed in an incubator to maintain constant temperature (25°C) for 14 hrs. At the end of the incubation period, the surface of the plates was imaged by microscopy, and subsequently the wells were re-suspended and phage concentration was determined by the plaque assay.

**Fig. 2.**
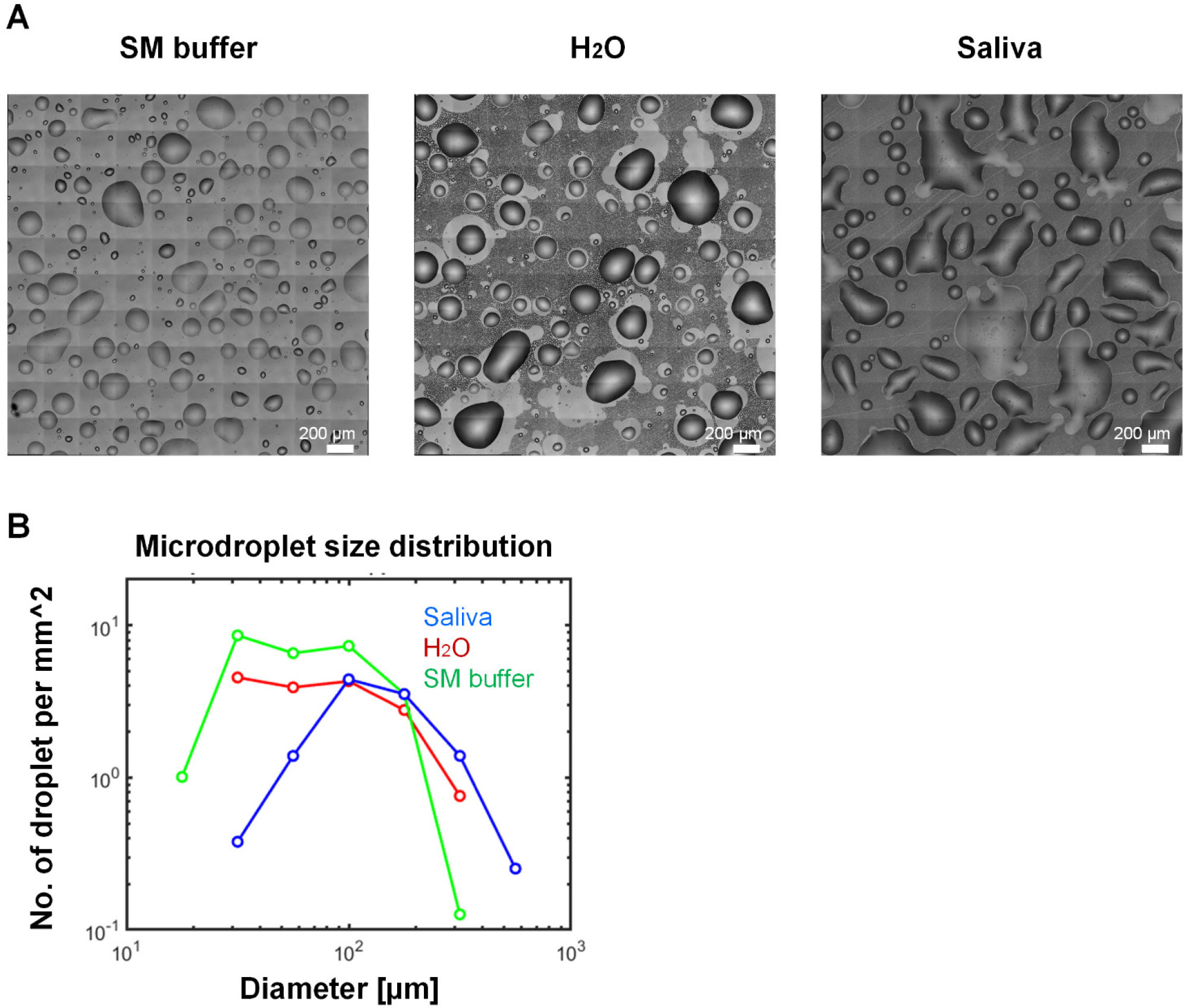
Representative microscopy images of sprayed SM, H_2_O, and human saliva on a glass surface. (**A**) typical shape and size of SM buffer, H_2_O, and saliva a few minutes after spray application onto a glass surface. (**B**) Droplet size distribution calculated based on the images shown in Panel A. Typical droplet size is between a few tens of microns to hundreds of microns in diameter, which is within the range of expelled saliva microdroplets that are expected to gravitate more rapidly toward surfaces (as opposed to aerosols, which remain suspended in the air for longer).

To better understand the microscale hydration state that viruses experience, we first examined the surfaces under the microscope 14 hrs post deposition. Representative images of the surface at t = 14 hrs are shown in Fig 3. Both the saliva and water microdroplets appeared completely dry at all tested RHs. As can be seen in Fig. 3, saliva microdroplets left a thin layer deposit (~3-5 μm thickness) of dry matter wherein natural bacterial flora could be seen as well as other aggregated substances, some of them are clearly salt crystals, but not all (somewhat similar to ^18^). Beads were dispersed fairly uniformly within these microdroplet deposits, as previously demonstrated for virions ^18^. While the beads are at least an order of magnitude larger than individual virions, their visualization can help us grasp the concentration of virions in droplets, and possibly their spatial distribution (assuming lack of virion aggregation, see Discussion). A back-of-the-envelope calculation estimated ~10 virions in a 100-μm droplet, which is consistent with the observed bead distribution (Fig. 3).

**Fig. 3.**
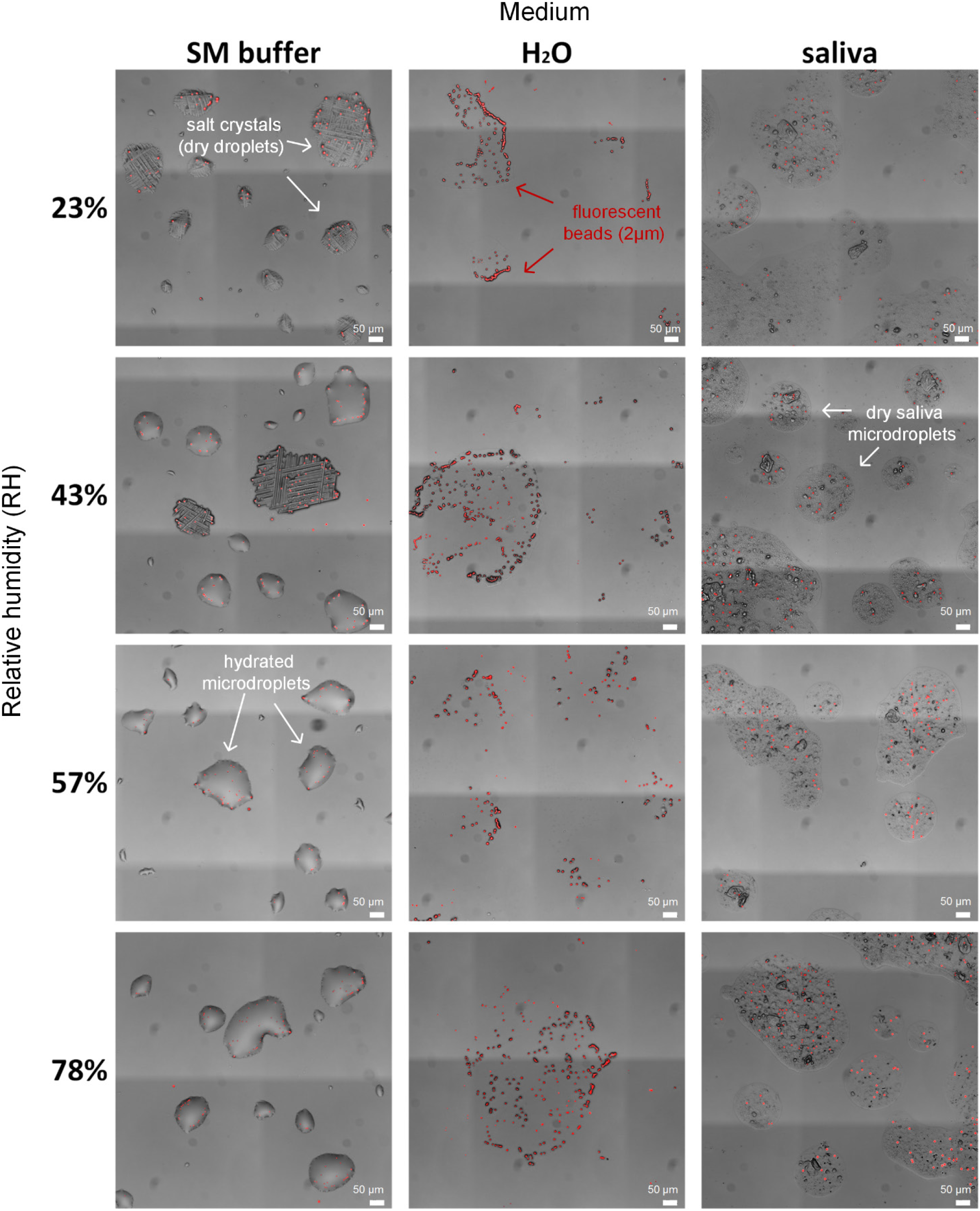
Microscopy images of microdroplets deposited on the glass-bottom well plates after 14 hrs incubation at a range of RH levels. SM buffer droplets remained hydrated at 57% RH and above, partially hydrated at 43% RH, and completely dry at 23% RH (medium salt crystallization can be seen in dried droplets). H_2_O and saliva droplets appeared to be dry under all tested RH conditions. Fluorescent micro-beads (2 μm) (red) were added at a concentration similar to the virus concentration, and thus may help to visualize the expected distribution of viruses within evaporated microdroplets.

Water microdroplets dried out at all RH levels. This observation is not surprising, as the water suspension contained very low concentrations of deliquescent solutes (e.g., salts). At RH = 78%, some tiny microdroplets (~ few μms in diameter) were observed around single beads or small bead clusters (Fig. 3). These tiny water microdroplets were likely retained due to strong capillary forces. In contrast, SM buffer microdroplets were hydrated at 57% RH and above (Fig. 3). Stable microdroplets of tens of μms in diameter could be clearly seen (Fig. 3). At RH = 43%, some of the microdroplets, but not all, dried out and salt crystals were observed. At RH = 23%, all microdroplets dried out and only salt crystals were seen (Fig. 3). These observations can be explained by the point of efflorescence of NaCl, the major solute in SM buffer: ~45-48% RH ^51^.

Next, we estimated virus survival under all tested conditions. The bottom surface of each well was suspended, and viral infectivity levels (PFU/ml) in the suspensions were evaluated by the plaque assay (Fig. 4 – see Methods). Virus viability in sprayed droplets (saliva, water, and SM buffer) was compared with that of suspended virus in bulk medium that was kept in sealed tubes throughout the duration of the experiment as a control. Strikingly, the highest survival of all tested viruses was found in evaporated saliva microdroplet deposits with less than 1.5 order of magnitude reduction in infectivity. A non-monotonous U-shaped survival as a function of RH was observed for Phi6 and PhiX174 with highest survival at 23% (even somewhat higher than the control). This is consistent with previous observations on superhydrophobic surfaces ^18^.

**Fig. 4.**
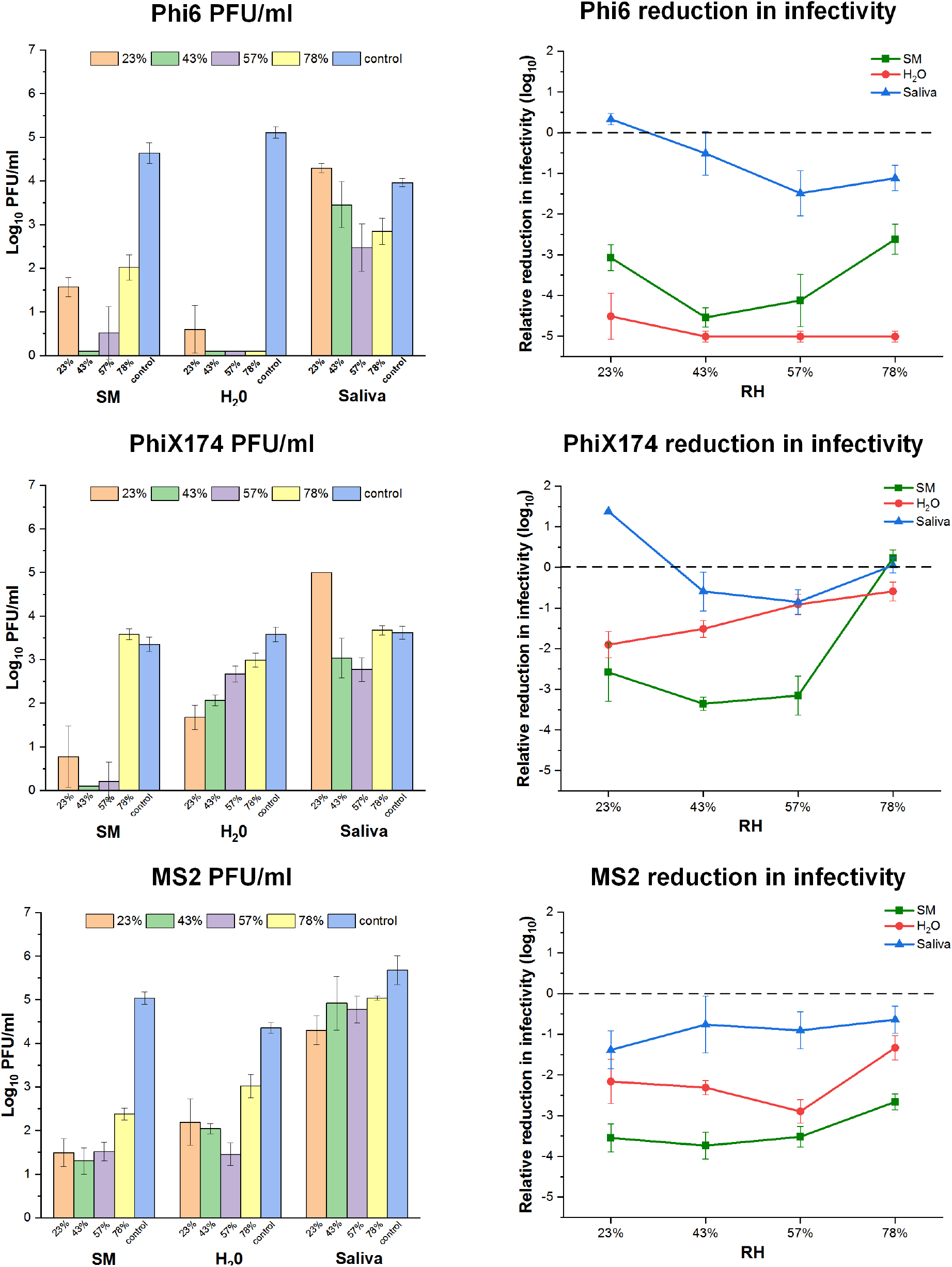
Viability of Phi6, PhiX174, and MS2 respectively as a function of RH and medium type on glass surface. **Left panel:** Bacteriophage titer concentration (PFU/ml) after 14 hrs incubation at 23%, 43%, 57%, and 78% RH at 25°C. A 10^5^-10^6^ titer of bacteriophage suspended in SM buffer, H_2_O, or human saliva was applied by spraying on a glass surface. A control sample was left suspended in the spraying device for the duration of the incubation period. Following the incubation period, phage concentration was determined by plaque assay (mean ± SD of four replicates). **Right panel:** Relative reduction of infectivity of Phi6, PhiX174, and MS2 in SM buffer (green), H_2_O (red) and saliva (blue) (mean ± SD of four replicates).

Similar to saliva, water microdroplets were dry at all RH levels. This appears to have a large impact on the enveloped virus Phi6, which showed more than 4 orders of magnitude reduction in infectivity (Fig. 4). In contrast, the two non-enveloped viruses showed lower reduction in infectivity (1-3 orders of magnitude, Fig. 4). While PhiX174 survival in evaporated water droplets increased with RH, hydration conditions did not show any noticeable change (Fig. 3). MS2 survival in water was lower than that in saliva, with a U-shaped RH dependence.

In the SM buffer, while microdroplets remained hydrated at 78% and 57% RH (Fig. 3), survival of all three viruses was lower than in saliva under the corresponding RH levels. The only exception was PhiX174, which exhibited similar survival levels in SM and saliva at 78% RH. Overall, Phi6, PhiX174, and MS2 in SM buffer deposits showed 3-4 orders of magnitude reduction in infectivity across the tested RH range, exhibiting the familiar U-shaped trend.

## Discussion

Aiming to obtain some mechanistic insights into factors that determine virus survival in microdroplets deposited on surfaces, we explored a wide range of RH, three types of solutions, and three model viruses (enveloped and non-enveloped). Combining surface microscopy imaging and infective virus enumeration, we were able to explore the link between the tested variables as manifested with respect to microscopic surface wetness and virus survival.

Our results indicate that RH and hydration conditions are not sufficient to explain virus survival in microdroplets deposited on surfaces. For example, the log reduction of Phi6 at 78% RH was ≈1, ≈3, and ≈5 when suspended in saliva, SM buffer, and H_2_O respectively. This is in contrast to high and comparable virus survival in bulk solution (control) of all tested media (Fig. 4). This implies that the physicochemical changes that characterize a microdroplet’s drying process have a pivotal implication for virus survival. Thus, the effect of RH on virus survival is context dependent. Likewise, SM droplets were hydrated at 78% RH and completely desiccated at 23% RH. Nonetheless, Phi6 survival under all of these widely differing hydration states was comparable (and all showing ~3-4 log reduction). We conclude that water availability and hydration status of a droplet cannot explain virus survival on its own. The observation that at a given RH, the microscopic hydration conditions of deposited droplets of various media can differ so widely (see along the rows of Fig. 3) suggests that RH does not directly affect virus stability and infectivity in drying microdroplets deposited on surfaces, but rather RH indirectly affects survival through its effect on physicochemical conditions at the scale that matters for viruses (~ μm).

Remarkably, both enveloped and non-enveloped viruses survive well in evaporated human saliva microdroplets deposited on inanimate surfaces at a wide range of RH levels. The observation that saliva droplets were completely dry even at 78% RH was somewhat surprising, as we expected saliva to effloresce at 40-50% RH ^51^. Human saliva is a complex fluid that somehow protects viruses in drying droplets and aerosols ^18,19^. A rough approximation estimates that in our experiments, droplets of ~100 microns in diameter contained around 10 virions. Thus, the mass of salts, proteins, and surfactants in these droplets are a few orders of magnitude higher than the total virion mass ^18^. As suggested by Marr et al., association between virion and saliva proteins (or other components) can protect the virions for prolonged periods (even days) in such dry saliva droplets on surfaces. In addition, the lower content of inorganic salts in saliva ^52^ than that in SM buffer may explain part of the large differences in virion survival between the two solutions. We remark that bacteriophage survival in human saliva is puzzling considering the fact that they are not expected to be selected by evolution for high survival in human saliva, unless they are infecting bacteria of the human microbiomes. We hypothesize that the structural stability of both human viruses and bacteriophages is an old, shared trait that is deeply entrenched in the evolution of viruses.

A second remarkable result is the very low survival (~5 log reduction) of Phi6 in the evaporated water microdroplets, regardless of RH. This is in stark contrast to the non-enveloped viruses that showed only moderate log reduction (~2) and moderate survival under high RH (78%). This result may indicate damage associated with the lipid membrane of the enveloped phage under these conditions. We note that in the bulk water, survival of Phi6 was high and comparable to that in SM buffer and saliva. A possible explanation therefor may be attributed to pH changes that occur during evaporation ^18^. The absence of protecting components such as salt crystals or saliva proteins may cause the damage.

This indicates that the physicochemical properties, experienced by the virions in the SM droplets from time of deposition through evaporation until reaching equilibrium, affect virus stability and viability. The dramatic increase in salt concentrations, and/or the time of exposure to high salt concentrations and reduced pH likely cause damage to the virus. This in turn suggests that the evaporation kinetics of suspended droplets on surfaces may affect the survival of viruses, in a similar manner as that suggested in aerosol droplets ^24^. Thus, complete dryness could in fact protect the virions from those high concentrations of dissolved salts and low pH ^5,19^.

Note that virus viability requires both physical stability of the virion and functionality, i.e., the ability to infect host cells and replicate. These in turn likely require stability of the capsid or envelope (depending on virus type), and that the relevant proteins (e.g., spike proteins), which play a role in attachment to host cells, remain functional. While the results of this study could not resolve between stability and infectivity, the differences between enveloped and non-enveloped viruses suggest that reduction in physical stability plays a role in virus viability.

Another less-understood issue is viral aggregation ^53,54^. Spontaneous aggregation of virions, for example as a response to changes in salt concentrations and pH ^46,53,55,^ may occur in drying microdroplets, and thus may play a role in virus survival. We speculate that with lipid-enveloped viruses suspended in water, aggregation of viruses imposed by hydrophobicity or other mechanisms, might play a vital role. If significant viral aggregation occurs, it has implications for interpretation of PFU/ml survival estimations (true for any virus survival study), and for the distribution of virions in drying droplets. Large virion aggregates (> tens of virions) might also affect microdroplets’ drying dynamics due to capillary pinning, as we have shown for bacteria ^34^.

The results of this study provide important insights concerning the COVID-19 pandemic. Although performed with a surrogate for SARS-CoV-2, it indicates, along with other studies, that SARS-CoV-2 survives well in evaporated saliva droplets on inanimate surfaces. Because SARS-Cov-2 transmission appears to depend upon viral load, it is likely that in indoor environments where infected individuals stay for long periods, viable viruses persist on fomites for days. Thus, as long as not proved otherwise, indirect transmission through inanimate surfaces – in particular those with prolonged and high contact – is not unlikely, and must be considered. A source of optimism may lie in the injurious impact of evaporating water droplets on the enveloped phage Phi6, an observation that should be confirmed with SARS-CoV-2.

## Acknowledgements

We thank Adi Stern for kindly providing the host *E. coli* strain of MS2. N. K. is supported by research grants from the James S. McDonnell Foundation (Studying Complex Systems Scholar Award, Grant #220020475) and from the Israel Science Foundation (ISF #1396/19).

## References

1 Organization, W. H. Modes of transmission of virus causing COVID-19: implications for IPC precaution recommendations: scientific brief, 27 March 2020. (World Health Organization, 2020).

2 Prather, K. A., Wang, C. C. & Schooley, R. T. Reducing transmission of SARS-CoV-2. Science (2020).

3 Dbouk, T. & Drikakis, D. On coughing and airborne droplet transmission to humans. Physics of Fluids 32, 053310, doi: https://doi.org/10.1063/5.0011960 (2020).

4 Duguid, J. The size and the duration of air-carriage of respiratory droplets and droplet-nuclei. Epidemiology & Infection 44, 471–479 (1946).

5 Yang, W. & Marr, L. C. Dynamics of airborne influenza A viruses indoors and dependence on humidity. PloS one 6 (2011).

6 Kampf, G. Potential role of inanimate surfaces for the spread of coronaviruses and their inactivation with disinfectant agents. Infection Prevention in Practice, 100044 (2020).

7 Otter, J. et al. Transmission of SARS and MERS coronaviruses and influenza virus in healthcare settings: the possible role of dry surface contamination. Journal of Hospital Infection 92, 235–250 (2016).

8 van Doremalen, N. et al. Aerosol and Surface Stability of SARS-CoV-2 as Compared with SARS-CoV-1 New England Journal of Medicine (2020).

9 Chin, A. et al. Stability of SARS-CoV-2 in different environmental conditions. medRxiv (2020).

10 Liu, Y. et al. Stability of SARS-CoV-2 on environmental surfaces and in human excreta. medRxiv, doi:https://doi.org/10.1101/2020.05.07.20094805 (2020).

11 Pan, Y., Zhang, D., Yang, P., Poon, L. L. & Wang, Q. Viral load of SARS-CoV-2 in clinical samples. The Lancet Infectious Diseases 20, 411–412 (2020).

12 Ye, G. et al. Environmental contamination of the SARS-CoV-2 in healthcare premises: An urgent call for protection for healthcare workers. medRxiv (2020).

13 Ong, S. W. X. et al. Air, surface environmental, and personal protective equipment contamination by severe acute respiratory syndrome coronavirus 2 (SARS-CoV-2) from a symptomatic patient. Jama 323, 1610–1612 (2020).

14 Chia, P. Y. et al. Detection of Air and Surface Contamination by Severe Acute Respiratory Syndrome Coronavirus 2 (SARS-CoV-2) in Hospital Rooms of Infected Patients. medRxiv, doi:https://doi.org/10.1101/2020.03.29.20046557 (2020).

15 Effros, R. M. et al. Dilution of respiratory solutes in exhaled condensates. American Journal of Respiratory and Critical Care Medicine 165, 663–669 (2002).

16 Larsson, B., Olivecrona, G. & Ericson, T. Lipids in human saliva. Archives of oral Biology 41, 105–110 (1996).

17 Schenkels, L. C., Veerman, E. C. & Nieuw Amerongen, A. V. Biochemical composition of human saliva in relation to other mucosal fluids. Critical reviews in oral biology & medicine 6, 161–175 (1995).

18 Vejerano, E. P. & Marr, L. C. Physico-chemical characteristics of evaporating respiratory fluid droplets. Journal of The Royal Society Interface 15, 20170939 (2018).

19 Santosh Kumar, S., Shao, S., Li, J., He, Z. & Hong, J. Droplet evaporation residue indicating SARS-COV-2 survivability on surfaces. arXiv, arXiv: 2005.12262 (2020).

20 Casanova, L. M., Jeon, S., Rutala, W. A., Weber, D. J. & Sobsey, M. D. Effects of air temperature and relative humidity on coronavirus survival on surfaces. Appl. Environ. Microbiol. 76, 2712–2717 (2010).

21 Van Doremalen, N., Bushmaker, T. & Munster, V. Stability of Middle East respiratory syndrome coronavirus (MERS-CoV) under different environmental conditions. Eurosurveillance 18, 20590 (2013).

22 Prussin, A. J. et al. Survival of the enveloped virus Phi6 in droplets as a function of relative humidity, absolute humidity, and temperature. Appl. Environ. Microbiol. 84, e00551–00518 (2018).

23 Shaman, J. & Kohn, M. Absolute humidity modulates influenza survival, transmission, and seasonality. Proceedings of the National Academy of Sciences 106, 3243–3248 (2009).

24 Lin, K. & Marr, L. C. Humidity-dependent decay of viruses, but not bacteria, in aerosols and droplets follows disinfection kinetics. Environmental Science & Technology 54, 1024–1032 (2019).

25 Kormuth, K. A. et al. Influenza virus infectivity is retained in aerosols and droplets independent of relative humidity. The Journal of infectious diseases 218, 739–747 (2018).

26 Yang, W. & Marr, L. C. Mechanisms by which ambient humidity may affect viruses in aerosols. Appl. Environ. Microbiol. 78, 6781–6788 (2012).

27 Yang, W., Elankumaran, S. & Marr, L. C. Relationship between humidity and influenza A viability in droplets and implications for influenza’s seasonality. PloS one 7 (2012).

28 Parker, E. R., Dunham, W. B. & Macneal, W. J. Resistance of the Melbourne Strain of Influenza Virus to Desiccation. Journal of Laboratory and Clinical Medicine 29, 37–32 (1944).

29 Thomas, Y. et al. Survival of influenza virus on banknotes. Appl. Environ. Microbiol. 74, 3002–3007 (2008).

30 Marr, L. C., Tang, J. W., Van Mullekom, J. & Lakdawala, S. S. Mechanistic insights into the effect of humidity on airborne influenza virus survival, transmission and incidence. Journal of the Royal Society Interface 16, 20180298 (2019).

31 Mauer, L. J. & Taylor, L. S. Water-solids interactions: deliquescence. Annual review of food science and technology 1, 41–63 (2010).

32 Wise, M. E., Martin, S. T., Russell, L. M. & Buseck, P. R. Water uptake by NaCl particles prior to deliquescence and the phase rule. Aerosol Science and Technology 42, 281–294 (2008).

33 Martin, S. T. Phase transitions of aqueous atmospheric particles. Chemical Reviews 100, 3403–3454 (2000).

34 Grinberg, M., Orevi, T., Steinberg, S. & Kashtan, N. Bacterial survival in microscopic surface wetness. eLife 8, e48508 (2019).

35 Rubasinghege, G. & Grassian, V. H. Role (s) of adsorbed water in the surface chemistry of environmental interfaces. Chemical Communications 49, 3071–3094 (2013).

36 Campbell, T. D. et al. Prebiotic condensation through wet–dry cycling regulated by deliquescence. Nature communications 10, 1–7 (2019).

37 Dallemagne, M. A., Huang, X. Y. & Eddingsaas, N. C. Variation in pH of model secondary organic aerosol during liquid–liquid phase separation. The Journal of Physical Chemistry A 120, 2868–2876 (2016).

38 Benbough, J. Some factors affecting the survival of airborne viruses. Journal of General Virology 10, 209–220 (1971).

39 Sun, Z.-p. et al. Stability of the COVID-19 virus under wet, dry and acidic conditions. medRxiv, doi:https://doi.org/10.1101/2020.04.09.20058875 (2020).

40 Turgeon, N., Toulouse, M.-J., Martel, B., Moineau, S. & Duchaine, C. Comparison of five bacteriophages as models for viral aerosol studies. Appl. Environ. Microbiol. 80, 4242–4250 (2014).

41 Kim, D.-K., Kim, S.-J. & Kang, D.-H. Inactivation modeling of human enteric virus surrogates, MS2, Qβ, and ΦX174, in water using UVC-LEDs, a novel disinfecting system. Food Research International 91, 115–123 (2017).

42 Rheinbaben, F., Schünemann, S., Gross, T. & Wolff, M. Transmission of viruses via contact in ahousehold setting: experiments using bacteriophage ϕX174 as a model virus. Journal of Hospital Infection 46, 61–66 (2000).

43 Aquino de Carvalho, N., Stachler, E. N., Cimabue, N. & Bibby, K. Evaluation of Phi6 persistence and suitability as an enveloped virus surrogate. Environmental science & technology 51, 8692–8700 (2017).

44 Ford, B. E. Pseudomonas Bacteriophage Phi6 as a Model for Virus Emergence. (2015).

45 Adcock, N. J. et al. The use of bacteriophages of the family Cystoviridae as surrogates for H5N1 highly pathogenic avian influenza viruses in persistence and inactivation studies. Journal of Environmental Science and Health, Part A 44, 1362–1366 (2009).

46 Langlet, J., Gaboriaud, F. & Gantzer, C. Effects of pH on plaque forming unit counts and aggregation of MS2 bacteriophage. Journal of Applied Microbiology 103, 1632–1638 (2007).

47 Bae, J. & Schwab, K. J. Evaluation of murine norovirus, feline calicivirus, poliovirus, and MS2 as surrogates for human norovirus in a model of viral persistence in surface water and groundwater. Appl. Environ. Microbiol. 74, 477–484 (2008).

48 Valegård, K., Liljas, L., Fridborg, K. & Unge, T. The three-dimensional structure of the bacterial virus MS2. Nature 345, 36–41 (1990).

49 McKenna, R. et al. Atomic structure of single-stranded DNA bacteriophage ϕX174 and its functional implications. Nature 355, 137–143 (1992).

50 Tang, J. W. The effect of environmental parameters on the survival of airborne infectious agents. Journal of the Royal Society Interface 6, S737–S746 (2009).

51 Posada, J., Redrow, J. & Celik, I. A mathematical model for predicting the viability of airborne viruses. Journal of virological methods 164, 88–95 (2010).

52 Pytko-Polonczyk, J., Jakubik, A., Przeklasa-Bierowiec, A. & Muszynska, B. Artificial saliva and its use in biological experiments. J Physiol Pharmacol 68, 807–813 (2017).

53 Gerba, C. P. & Betancourt, W. Q. Viral aggregation: impact on virus behavior in the environment. Environmental science & technology 51, 7318–7325 (2017).

54 Andreu-Moreno, I. & Sanjuán, R. Collective Viral Spread Mediated by Virion Aggregates Promotes the Evolution of Defective Interfering Particles. mBio 11 (2020).

55 Floyd, R. & Sharp, D. Viral aggregation: effects of salts on the aggregation of poliovirus and reovirus at low pH. Appl. Environ. Microbiol. 35, 1084–1094 (1978).

